# Human magnetic field registration capability using low-field magnetic sensors

**DOI:** 10.1101/2021.06.04.447144

**Authors:** Natalija Bricina, Jurijs Dehtjars, Ksenija Jašina, Hermanis Sorokins

## Abstract

The aim of the study was to find out whether it is possible to detect the human magnetic field without shielding using sensors. For this, an experiment “the possibility of detecting the human magnetic field using sensors” was conducted. During the experiment it was found that with the help of a low-field magnetic detector it is possible to register a person’s magnetic field, which is changing if there is a change in a heart rate.

During the study, a device for recording the magnetic field of a human was assembled, a methodology for obtaining data was developed, a description of the materials and equipment used was provided. In accordance with the described methods of data collection, measurements of human magnetic field with and without load were made. The data obtained during the study were processed and the results were displayed. Based on the obtained results, an analysis was carried out, conclusions were drawn and recommendations for subsequent studies were indicated. As a result, the possibility of detecting a human’s magnetic field using sensors was first studied.

## Introduction

It is known that a living organism generates magnetic fields. There are three sources of magnetic fields in the body, one of which is muscle and nerve cell activity, and the other is ferromagnetic particles that are brought into the body. The most active source of the magnetic field is the internal organs. The heart is the most powerful source of the body’s magnetic field. However, the heart’s magnetic field is much smaller than the Earth’s magnetic field, and it is known that as you move away from the heart, the intensity of the magnetic field decreases. [3]

There are currently several methods for recording a human magnetic field, including magnetocardiography. The main disadvantage of the method is that in order to obtain measurements, the patient must be in a specially isolated room where the external magnetic field does not flow. To obtain more accurate measurements, it is necessary to ensure complete immobility of the patient, as well as to remove devices that may generate a magnetic field. [4]

However, detectors have emerged that can detect weak magnetic fields. Therefore, can be investigated whether the use of these detectors can help to detect a human magnetic field using sensors that would be performed in an open space.

It is hypothesized that with the help of commercial detectors it is possible to record a human magnetic field in an open space, separating it from the total received magnetic field signal with a connection to the heart rate. This means that the detector is sensitive enough to detect particularly weak magnetic fields generated by the heart and no shielding is required.

The aim of the study is to find out whether it is possible to determine the human magnetic field using sensors. An experiment with a commercial detector was carried out for the first time. The results of the study can be used to develop a method for recording the human magnetic field using sensors, as well as to design a measuring device.

## Methods

### Human magnetic field registration with sensors

Since the human circulatory system is the source of the biocurrent and magnetic field, measurements were carried out by changing the blood flow and recording the ECG and magnetic field. The change in blood flow makes it possible to determine a human magnetic field. To determine human magnetic field detection capability with sensors, the triple axis compass magnetometer sensor module HW-127 HMC5883L (made by Honeywell), designed for low field magnetic sensing, was used. The minimal output rate of the detector (HMC5883L) is 0.75 Hz; this allows to detect the magnetic field generated by human heart with a heart rate starting from 45 bpm. [1,2] The detector (HMC5883L) was connected to the Arduino MEGA 2560 board using wires. The low-field magnetic detector (HMC5883L) was attached on the top of the registration board that was made from the cardboard (see Figure 1.). The detector (HMC5883L) was tightly fixed to the registration board with wool thread. It is important not to use any metal parts so that they do not create an additional background magnetic field. ECG recording was performed using a built-in signal conditioning unit (AD8232, manufactured by SparkFun).

**Fig.1.**
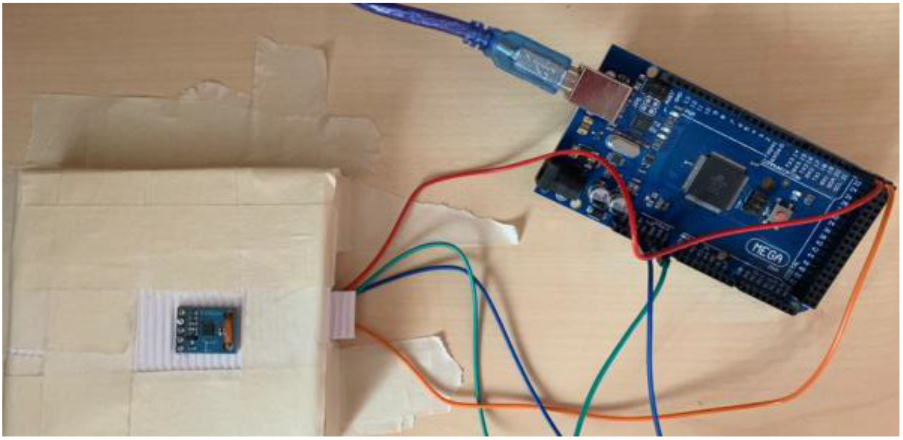
The HMC5883L detector location on the cardboard attached to the Arduino MEGA 2560 board.

The received signal from the detector (HMC5883L) was divided into intervals, each of which is 5000 ms. Each interval with a compressed hand was compared to the interval when the hand was decompressed. The experiment was repeated after the subject did physical exercise.

Experimentally the upper arm was compressed as hard as possible, so that the blood vessels did not allow blood to pass through. The blood flow was changed with a cuff, which, when compressed, will significantly reduce the blood flow in the upper arm. A cuff with a bulb is used, as it is possible to quickly and easily control the pressure in the cuff, as well as to release air quickly from the cuff using valve. A cuff is put on the upper arm of the right hand. The arm with the cuff is placed on the detector (HMC5883L). The inner part of the wrist area of the hand touches the detector (HMC5883L). Any movement can affect the result, so the elbow is supported, and hand is fully relaxed.

To perform the experiment with the ECG, electrodes are tightly attached to the skin surface. The electrodes were attached to the three points on the body. Where R and L electrodes were attached to the skin area under the clavicle on the right and left side respectively. The electrode COM was attached to the skin surface above the right iliac crest.

In order to determine the relation of the obtained magnetic field signal to the heart rate, an electrocardiogram registration was recorded together with the magnetic field. By combining two signals, the time values for the ECG R-peak, as it shows the electric impulse that is generated in heart and the maximum of the magnetic field correlation coefficient following the R-peak signal were recorded (see Figure 2.). Then the ECG and magnetic field time values were compared manually, peak values were found. The experiment with load was repeated.

**Fig.2.**
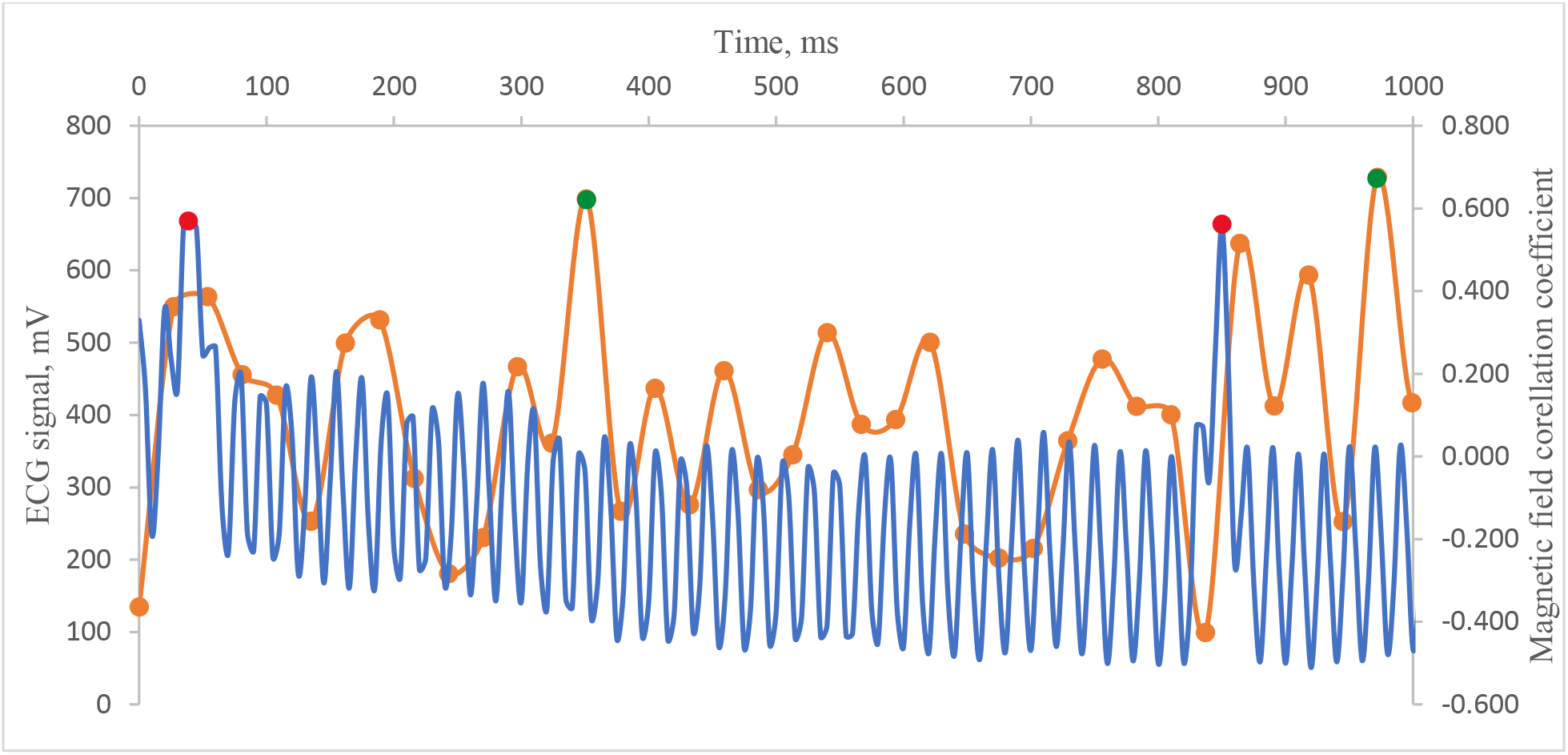
Detection of the magnetic field correlation coefficient maximum peak following the ECG R-edge signal. *Blue line* – ECG signal; *Orange line –* magnetic field correlation coefficient; *Red point* – selected ECG R-edge; *Green point –* selected magnetic field correlation coefficient maximum following the ECG R-edge signal.

At the start of the experiment, the cuff is compressed to 280-300 mmHg to ensure complete cessation of blood flow to the hand. It was made sure that there is some discomfort in the upper arm when the cuff is compressed. The right hand with the cuff is placed over the detector. After that, the magnetic field of the inner side of the wrist was recorded. With the compressed cuff, the recording of the signal was provided for 30 seconds to do not discomfort subject for too long. When 30 seconds have elapsed, air is rapidly expelled from the cuff and data is recorded with an open blood flow for another 30 seconds. For air to expel from the cuff at least 2 seconds are needed. The cuff is compressed again within the last 5 seconds to ensure that the cuff is already compressed the next minute when the second cycle begins. The measurements are performed in five cycles, which means that the total time for one experiment is 5 minutes. The body was allowed to rest for at least 1.5 hours before the experiment was repeated with load. Before taking measurements, 50 squats were made as fast as possible. When the heart rate was increased, measurements were taken immediately. To reduce the preparation time for the experiment, it is recommended to perform squats already with tightly attached electrodes.

To check whether the ECG affects the data the experiment was repeated with the same algorithm without registering a cardiogram. Before the experiment started, the pulse was measured with a blood pressure monitor. Afterwards, the experiment with load of 50 squats was repeated.

## Results

### Human magnetic field with sensors

The very first experimental results showed that there is a difference between a magnetic field signal with and without blood flow (see Figure 3.). The appearance of the signal with load and without is also different (see Figure 4.).

**Fig.3.**
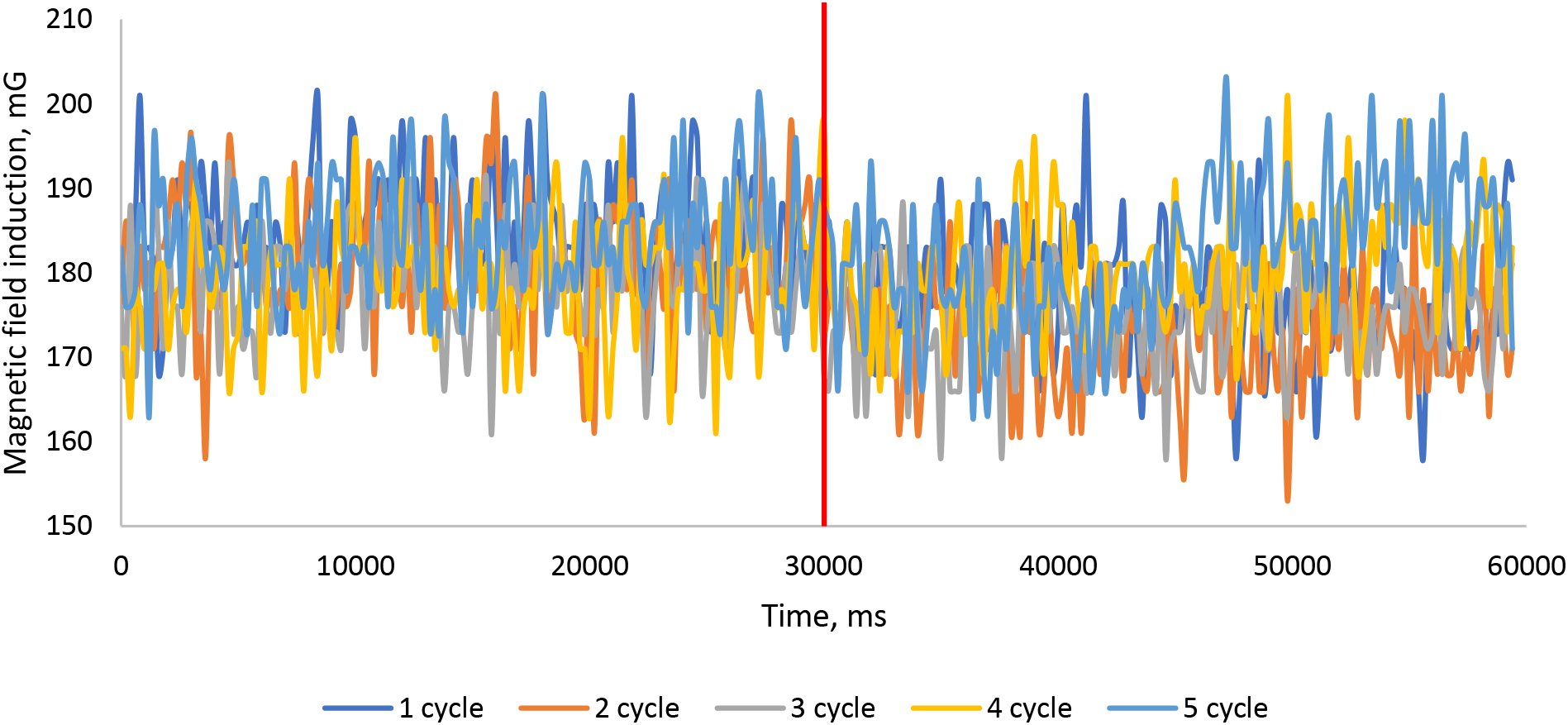
Magnetic field induction for five cycles, without load, where the left side of the graph shows signal when the hand is compressed; the right side of the graph shows signal when the hand is decompressed. *Red vertical line –* moment of time when air was released from the cuff. Magnetic field induction values are provided in y-direction.

**Fig.4.**
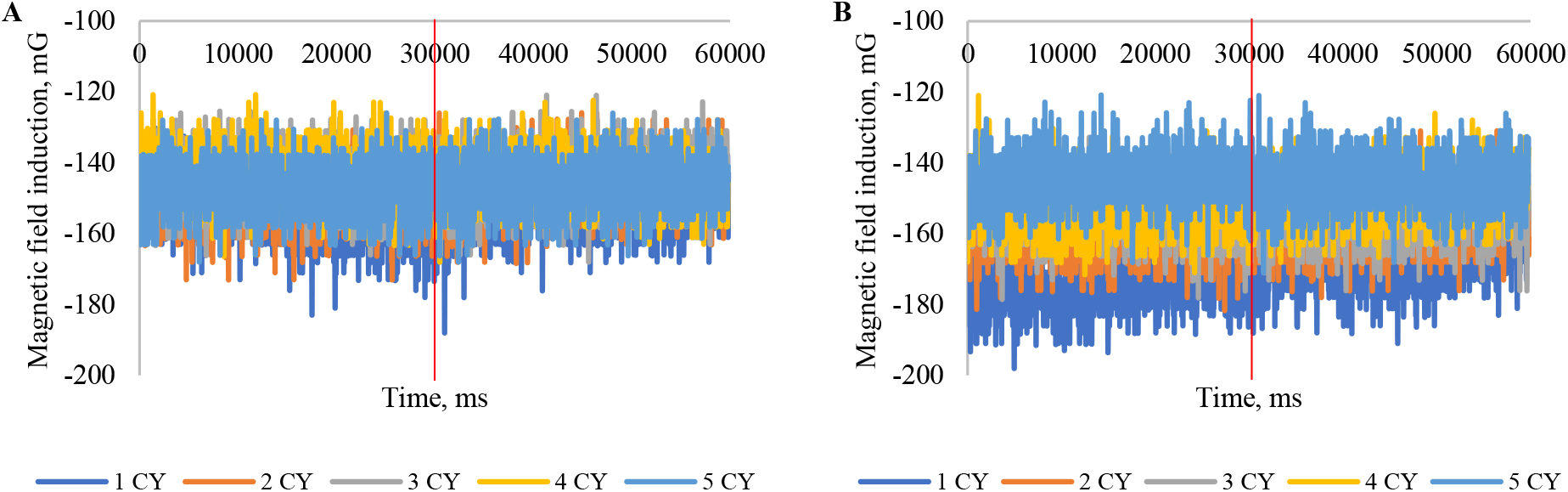
Magnetic field induction for five cycles, **a** with load heart rate is 136 bpm, **b** without load heart rate is 70 bpm, where the left side of the graph shows signal when the hand is compressed; the right side of the graph shows signal when the hand is decompressed. *Red vertical line –* moment of time when air was released from the cuff. Magnetic field induction values are provided in y-direction. The amplitude of magnetic field induction shifts when load is not applied.

In the case when load is not applied (heart rate is 70 beats per minute) the starting point of the signal changes every cycle. There is an amplitude shift of the signal. However, when load is applied (heart rate is 136 beats per minute), the starting point remains the same.

The comparison between phase was made; it shows that there was a difference in peak number on the time interval between experiment with and without load.

As for the experiments with ECG, the time difference between R-peak and following magnetic field peak on each interval of 5000 ms was calculated. The time difference in experiment without load is higher than with load in general. However, it changes in pulses and has fluctuations (see Table 1.) The time difference when load is applied is more constant, but there are peaks on intervals when blood flows (see Table 2.).

**Table 1.** Time difference (ms) between ECG R-peak and magnetic field following peak signal; without load

| Time interval (s) | Mean | SD |
| --- | --- | --- |
| 0-5 | 221 | 116 |
| 5-10 | 283 | 87 |
| 10-15 | 110 | 9 |
| 15-20 | 98 | 30 |
| 20-25 | 174 | 78 |
| 25-30 | 133 | 75 |
| 30-35 | 117 | 31 |
| 35-40 | 119 | 31 |
| 40-45 | 212 | 184 |
| 45-50 | 55 | 36 |
| 50-55 | 95 | 67 |
| 55-60 | 74 | 45 |

**Table 2.** Time difference (ms) between ECG R-peak and magnetic field following peak signal; with load

| Time interval (s) | Mean | SD |
| --- | --- | --- |
| 0-5 | 69 | 36 |
| 5-10 | 80 | 46 |
| 10-15 | 99 | 68 |
| 15-20 | 106 | 34 |
| 20-25 | 88 | 30 |
| 25-30 | 71 | 38 |
| 30-35 | 81 | 52 |
| 35-40 | 81 | 49 |
| 40-45 | 143 | 31 |
| 45-50 | 158 | 113 |
| 50-55 | 87 | 26 |
| 55-60 | 149 | 43 |

When the load is not applied, the time difference peak reaches its value just when air is released from the cuff. This means that the time difference between the electrocardiogram signal and the magnetic field signal increases within the time when the arm is compressed. Thus, the longer the arm is compressed, the later the magnetic field signal that follows the electric field signal is observed. Analyzing the part of the graph where there is no air in the cuff it can be observed that the time difference starts to decrease gradually. The time difference changes pulsatile with both compressed and relaxed hand.

In the experiment with load, the time difference changes are also pulsating, but the peak value of time is reached later after the air from the cuff is released. This could indicate that the body’s response to the blood flow stop, if a load was applied is different from a no-load response. (see Figure 5.).

**Fig.5.**
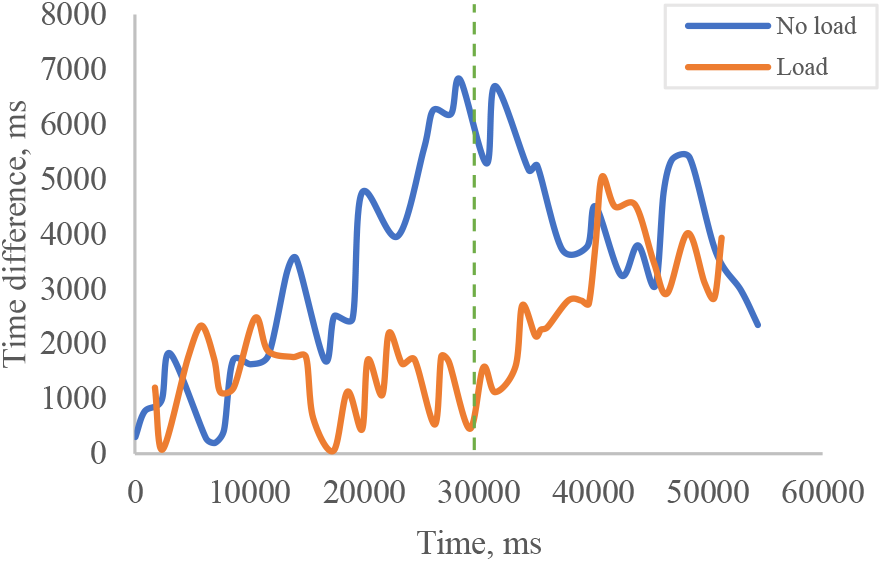
Plot of time difference between ECG and magnetic field when time values are compared, and peaks found. *Blue line* represents time difference when load is not applied; *Orange line* represents time difference when load is applied; *Green line* represents the moment when air is released from the cuff.

## Discussion

To the best of our knowledge, our study is the first to investigate how human magnetic field using sensors can be measured in open space with commercial low-field magnetic detectors. Our main findings are that the low-field magnetic sensors can detect human magnetic field. Further, the magnetic field depends on the blood flow and the biocurrent that is produced by the organism. This assumption is underpinned by the feedback of the organism, when the heart rate is changed.

These findings were surprising, since the human body generated alternating magnetic fields are in the range of 0.1-0.2 mkT [1]

## Conclusion

The aim of the experiment was to find out whether it is possible to detect the human magnetic field using sensors, an experiment with a commercial detector was carried out for the first time. It was decided to change the blood flow and a heart rate to observe changes in the magnetic field.

Experiment shows that the magnetic field signal amplitude shifts when heart rate is changed (70 bpm and 136 bpm). When the heart rate is 70 bpm the magnetic field amplitude shift is 25±11 mG and when the heart rate is 136 bpm the shift is 6±10 mG. Furthermore, the time difference between the ECG and the magnetic field correlation coefficient on the time intervals is different when heart rate is changed. The difference in time difference between load and non-load experiment on the time interval from 5-10 seconds was the largest. Time difference in experiment without load is 283±87 ms and 80±46 ms when load is applied. However, the smallest difference in time difference between two experiments was observed on the time interval from 50-55 ms: experiment without load 95±67 ms and with load 87±26 ms. As a result, it is possible to detect human magnetic field with sensors.

Our results may serve to stimulate future investigations into the magnetic field detection capability on different age groups.

## Notes

### Competing Interest Statement

The authors have declared no competing interest.

